# Testing for the genomic footprint of conflict between life stages in an angiosperm and moss species

**DOI:** 10.1101/2024.10.04.613734

**Authors:** Meng Yuan, Leslie M. Kollar, Bianca M. Sacchi, Sarah B. Carey, Baharul I. Choudhury, Teresa Jones, Jane Grimwood, Spencer C. H. Barrett, Stuart F. McDaniel, Stephen I. Wright, John R. Stinchcombe

## Abstract

The maintenance of genetic variation by balancing selection is of considerable interest to evolutionary biologists. An important but understudied potential driver of balancing selection is antagonistic pleiotropy between diploid and haploid stages of the plant life cycle. Despite sharing a common genome, sporophytes (2*n*) and gametophytes (*n*) may undergo differential or even opposing selection. Theoretical work suggests the antagonistic pleiotropy between life stages can generate balancing selection and maintain genetic variation. Despite the potential for far-reaching consequences of gametophytic selection, empirical tests of its pleiotropic effects (neutral, synergistic, or antagonistic) on sporophytes are generally lacking. Here, we examined the population genomic signals of selection across life stages in the angiosperm *Rumex hastatulus* and the moss *Ceratodon purpureus*. We compared gene expression among life stages and between sexes, combined with neutral diversity statistics and the analysis of the distribution of fitness effects. In contrast to what would be predicted under balancing selection due to antagonistic pleiotropy, we found that unbiased genes between life stages were under stronger purifying selection, likely explained by a predominance of synergistic pleiotropy between life stages and strong purifying selection on broadly expressed genes. In addition, we found that 30% of candidate genes under balancing selection in *R. hastatulus* were located within inversion polymorphisms. Our findings provide novel insights into the genome-wide characteristics and consequences of plant gametophytic selection.

**Significance:** The genome-wide importance of balancing selection has been a key question in evolutionary biology. Despite theoretical support for the potential of balancing selection generated by antagonistic pleiotropy between plant life stages, to our knowledge no empirical studies have systematically investigated this on a genome-wide scale. Our results revealed genome-wide patterns more consistent with synergistic pleiotropy, with gametophyte-biased genes more likely to be under relaxed purifying selection in both the angiosperm and moss species we investigated. In addition, our study suggests gene expression level and breadth has a greater effect on selection efficacy than differences between ploidy levels in different life stages.

## Introduction

Understanding how genetic variation is maintained for traits under selection remains a key open question in evolutionary biology. Balancing selection maintains genetic variation in a population when different alleles are favored in different contexts, e.g., antagonistic selection between sexes or life history, hereafter life stages (Kidwell et al. 1977; Immler et al. 2012). Despite persistent theoretical and empirical interests, we still lack empirical tests of the genome-wide importance of balancing selection. The increasing availability of genomic data helps us to understand how different forms of balancing selection shape genetic variation, and also to identify candidate regions under balancing selection (Bitarello et al. 2023; Ruzicka et al. 2025). Here, we focus on testing for genome-wide signals of balancing selection generated by antagonistic pleiotropy between life stages using transcriptomic and population genomic data of the angiosperm *Rumex hastatulus* and the moss *Ceratodon purpureus*.

The alternation of life cycles between diploid and haploid phases in sexually reproducing eukaryotes allows for natural selection to occur during both phases (Mable and Otto 1998). In land plants, the haploid phase (gametophyte) and diploid phase (sporophyte) have varying degrees of complexity in the different major clades of land plants. For example, angiosperms generally have the most elaborated sporophytes and the most reduced gametophytes; their gametophytes (i.e., ovules and pollen) are smaller and ephemeral, have fewer cells, and are dependent on sporophytes. Contrastingly, the haploid gametophytes in bryophytes (mosses, liverworts, and hornworts) are multicellular and often perennial. Differential or even opposing selection can result from the differences in genetic, cellular and organismal levels between these two life stages, creating genomic conflicts (Qiu et al. 2012; reviewed in Immler 2019). How selection on gametophytes (gametophytic selection) affects sporophytes is a key unresolved question in plant evolutionary genetics.

The evolutionary consequences of gametophytic selection have been examined both theoretically and empirically (Haldane 1932; Otto et al. 2015; Immler and Otto 2018; summarized in Immler 2019, Beaudry et al. 2020). The haploid nature of the gametophytic phase is predicted to increase the efficacy of selection due to the lack of dominance (i.e., the masking hypothesis), allowing for more efficient removal of recessive deleterious mutations (i.e., purging) and fixation of recessive beneficial mutations (Crow and Kimura 1965; Kondrashov and Crow 1991; Gerstein and Otto 2009). Consistent with this prediction, some studies have found stronger purifying and positive selection in gametophyte-specific than sporophyte-specific genes (Arunkumar et al. 2013; Cervantes et al. 2023; but see Harrison et al. 2019). Stronger selection on gametophyte-expressed genes can also be driven by gametophytic competition (sexual selection) in angiosperms (Arunkumar et al. 2013; Gossmann et al. 2014; Gutiérrez-Valencia et al. 2022). Additionally, gene expression breadth can be a stronger predictor of selection efficacy than the masking effect in angiosperms and bryophytes (Szövényi et al. 2013).

The common genome shared between life stages provides an opportunity to test how gametophytic selection affects the evolution of the sporophyte stage due to pleiotropy between life stages. Plants share a substantial overlap in gene expression between life stages based on both microarray and mRNA sequencing studies (summarized in Beaudry et al. 2020). In ferns and bryophytes where the gametophytes are photosynthetic and exhibit indeterminate growth, gene expression overlap may exceed that in angiosperms, e.g., estimates are ∼60% in the angiosperm *Arabidopsis thaliana* (Walbot and Evans 2003; Borg et al. 2009), 97.7% in the fern *Polypodium amorphum* (Sigel et al. 2018), and 85% in the bryophyte *Physcomitrium patens* (Ortiz-Ramírez et al. 2016). Previous studies on gametophytic selection have compared selective forces in gametophyte *-vs.-* sporophyte-specific genes to test for the effects of haploid expression on the distribution of selection coefficients (Arunkumar et al. 2013; Gossmann et al. 2014; Harrison et al. 2019; Gutiérrez-Valencia et al. 2021; Cervantes et al. 2023). However, the patterns of genetic diversity and selection pressures of genes with overlapping expression, which are more likely to be pleiotropic due to the greater potential for competing selection pressures across life stages (Immler et al. 2012; Peters and Weis 2018), have not yet been specifically examined.

In theory, antagonistic pleiotropy between life stages, i.e., intralocus conflict when alleles of a gene have opposite fitness effects between life stages, can cause balancing selection and maintain genetic variation under some conditions (Immler et al. 2012; Peters and Weis 2018). Despite the theoretical support, empirical evidence of antagonistic pleiotropy in angiosperms is limited to a few studies suggesting sexual conflict across life stages, e.g., between male gametophyte and female sporophyte during pollen competition (e.g., Travers and Mazer 2001; Lankinen and Kiboi 2007), and between female gametophyte and male sporophyte in a female meiotic drive system (Fishman and Saunders 2008). In bryophytes where sporophytes are dependent only on female gametophytes, parent offspring conflict may occur (Haig and Wilczek 2006; Johnson and Shaw 2016; Shortlidge et al. 2021), but the mechanism and genetic architecture underlying any such conflict remain unclear. Testing the genome-wide prevalence of antagonistic pleiotropy between life stages and its potential to maintain genetic variation is therefore key to understanding the genetic architecture of conflict between plant life stages.

Given the potential for antagonistic pleiotropy between the life stages for genes with overlapping expression, we predicted that if these effects are widespread, we should see population genetic signals of balancing selection more commonly in genes with overlapping expression (Immler et al. 2012; Peters and Weis 2018). An analogous finding is that genes with intermediate rather than extreme sex bias between sexes show patterns of genetic diversity consistent with balancing selection, as expected if extreme expression bias reflects resolved rather than ongoing conflict (e.g., Cheng and Kirkpatrick 2016; Wright et al. 2018; Sayadi et al. 2019; summarized in Ruzika et al. 2020). Previous studies on sexual conflict and sex-biased expression studies primarily relied on diversity statistics. Recently developed model-based approaches offer an additional way to detect balancing selection (Bitarello et al. 2023) and assess enrichment of biased genes between life stages.

We examined gene expression and genetic diversity patterns in the gametophytic and sporophytic life stages of two plant species: the angiosperm *Rumex hastatulus* (Polygonaceae) and the bryophyte *Ceratodon purpureus* (Ditrichaceae). The annual *R. hastatulus* is dioecious, obligately outcrossing, and wind-pollinated with an XY sex chromosome system (Smith 1964; Rifkin et al. 2021). The uniovulate flowers and the large quantity of pollen commonly produced in wind-pollinated angiosperms suggests high potential for gametophytic selection during pollen competition (Friedman and Barrett 2009; Field et al. 2012; Immler 2019). In the dioicous moss *C. purpureus*, the female and male gametophytes possess the U or V sex chromosome, respectively (Carey et al. 2021a), they produce gametes that fuse to form a diploid zygote (i.e., sporophyte, unsexed); the sporophyte matures and ultimately undergoes meiosis, but its entire development occurs on the female gametophyte plant, providing multiple opportunities for antagonistic pleiotropy (Carey et al. 2021b; McDaniel 2023). The contrasts in the length and complexity of the gametophytic stage in these two species allow us to evaluate the evolutionary consequences of biphasic plant life cycles in a broader context.

We first compared gene expression in gametophytic and sporophytic tissues of *R. hastatulus* and *C. purpureus,* finding a larger overlap in gene expression and a smaller proportion of biased genes between life stages in *C. purpureus* than *R. hastatulus*. We then examined the population genetic signals of balancing selection due to antagonistic pleiotropy between life stages by examining nucleotide diversity and Tajima’s *D* across varying degrees of expression bias between life stages. In contrast with our predictions for genome-wide patterns of antagonistic pleiotropy, our results show elevated nucleotide diversity and Tajima’s *D* in highly gametophyte-biased genes in both *R. hastatulus* and *C. purpureus*. We conclude that expression levels and breadth were a stronger predictor of selection efficacy than potential competing selective pressures between life stages. Lastly, we identified in both species genomic regions containing hundreds of candidate genes under balancing selection using a composite likelihood ratio test (Cheng and DeGiorgio 2019, 2020, 2022) and examined the biological functions and differential gene expression patterns of these candidate genes.

## Results

### Gene expression in gametophytes and sporophytes

We compared gene expression in gametophyte and sporophyte tissues of *R. hastatulus* and *C. purpureus*. In *R. hastatulus*, we sequenced 75 mature pollen samples and 77 male leaf samples representing the gametophyte and sporophyte stages, respectively (Table S1). We used only male leaf tissue for assessing sporophytic expression in *R. hastatulus* to avoid confounding effects from potential sexual conflict between leaves of males and females. In *C. purpureus*, we compared expression between 21 sporophyte and 36 gametophyte samples (whole plant, described in Carey et al. 2021a, Table S3, see Materials and Methods). In both species, there is a large overlap in gene expression between the life stages (Table 1). In *R. hastatulus*, we identified 15,958 gametophyte-expressed genes, consisting of 52% of all annotated genes across the genome, and 89% of genes that were expressed in either life stage. In *C. purpureus*, we identified 24,097 gametophyte-expressed genes, consisting of 69% of all annotated genes and 93% of genes that were expressed in either life stage. The overlap in gene expression between life stages (89% in *R. hastatulus*, and 93% in *C. purpureus*) sets up a strong potential for pleiotropic effects of mutations between life stages.

**Table 1.** Summary of samples used and gene expression analyses in *R. hastatulus* and *C. purpureus*. *P*-adj: Benjamini-Hochberg adjusted *p*-value with a false discovery rate of 0.1, FC: fold change in expression, i.e., the ratio of expression level between the two tissues compared. All percentages are based on comparison to the number of expressed genes.

|  | Genome size | Sample size of RNA samples | Number of annotated genes | Number of expressed genes | Number of gametophyte-expressed genes | Number of biased genes (*p*<sub>adj</sub> < 0.1, FC > 2) |
| --- | --- | --- | --- | --- | --- | --- |
| *Rumex hastatulus* | 1.7 Gb (2n) | 75 (n),<br>77 (2n) | 30,641 (2n) | 17,984 | 15,958 (89%) | 5,635 (n, 31%)<br>8,653 (2n, 48%) |
| *Ceratodon purpureus* | 358 Mb (n) | 36 (n),<br>21 (2n) | 34,392 (2n) | 25,880 | 24,097 (93%) | 6,446 (n, 25%)<br>5,202 (2n, 20%) |

We first identified genes that were differentially expressed (DE) between life stages and exclusively expressed in one life stage (Table S4). There was a slightly higher percentage of overlapping gene expression and a lower percentage of differentially expressed genes between life stages in *C. purpureus* than *R. hastatulus* (Table 1). In *R. hastatulus*, we identified 5,635 gametophyte-biased and 8,653 sporophyte-biased genes; similarly in *C. purpureus*, there were 6,446 gametophyte-biased and 5,202 sporophyte-biased genes (Table 1). Among significant DE genes, we filtered for genes in the top 10% quantile of expression bias, allowing us to compare genes with different tissue specificities. Our filtering process resulted in similar numbers of gametophyte- and sporophyte-specific genes: 1,711 gametophyte-specific and 1,711 sporophyte-specific genes in *R. hastatulus*, and 1,886 gametophyte-specific and 1,867 sporophyte-specific genes in *C. purpureus*. Gene ontology (GO) enrichment analyses indicated very different functional enrichment for biased and specific genes between life stages (Table S5, S6), highlighting distinctive expression profiles across life stages in both species. For example, in *R. hastatulus*, gametophyte-specific genes had protein phosphorylation functions (GO:0006468) which are important during pollen-tube growth (Klodová and Fíla 2021) and sporophyte-specific genes showed photosynthetic functions (GO:0015979).

### Nucleotide diversity and expression bias between life stages

Population genomic analyses can be used to provide an indirect test for pleiotropic effects across life stages. We calculated nucleotide diversity at 4-fold (*π*_s_) and 0-fold (*π*_n_) sites for each gene and looked at the linear relation between *π*_s_ or *π*_n_ and the direction and degree of expression bias between life stages (log2FoldChange) across genes. The linear regression of *π*_s_ or *π*_n_ on the direction and degree of expression bias between life stages (log2FoldChange and its absolute value, where positive values indicate gametophyte bias and negative values indicate sporophyte-bias) showed a significant but weak positive relationship, with near-zero slopes in both species (Figure S1, S2). We next divided the genes into eight bins based on the direction and degree of expression bias between life stages and calculated the weighted mean of *π*_s_ and *π*_n_ for each bin, weighted by the number of sites per gene. In both species, highly gametophyte-biased genes exhibited the highest levels of *π*_s_, *π*_n_ and *π*_n_/*π*_s_ ratio (Figure 1, Table S7). This result contrasts with our expectation that genes with overlapping and intermediate bias in expression should show the highest diversity under a model of widespread antagonistic pleiotropy. In both species, *π*_s_ and *π*_n_ increase with the degree of expression bias between life stages in both directions (less so in *R. hastatulus*), but values of *π*_s_ and *π*_n_ were greatest for the highly gametophytic-biased genes (Figure 1). In *R. hastatulus*, the increase in *π*_n_ with the degree of expression bias was less symmetrical than in *π*_s_, with sporophyte-biased genes having lower *π*_n_ than genes with overlapping expression (Figure 1a, 1b). Our results were robust to our choice of weighting *π*_s_ and *π*_n_ by the number of sites per gene, as when we estimated the mean and standard errors without weighting by the number of sites, the patterns remained the same (Figure S3).

**Figure 1.**
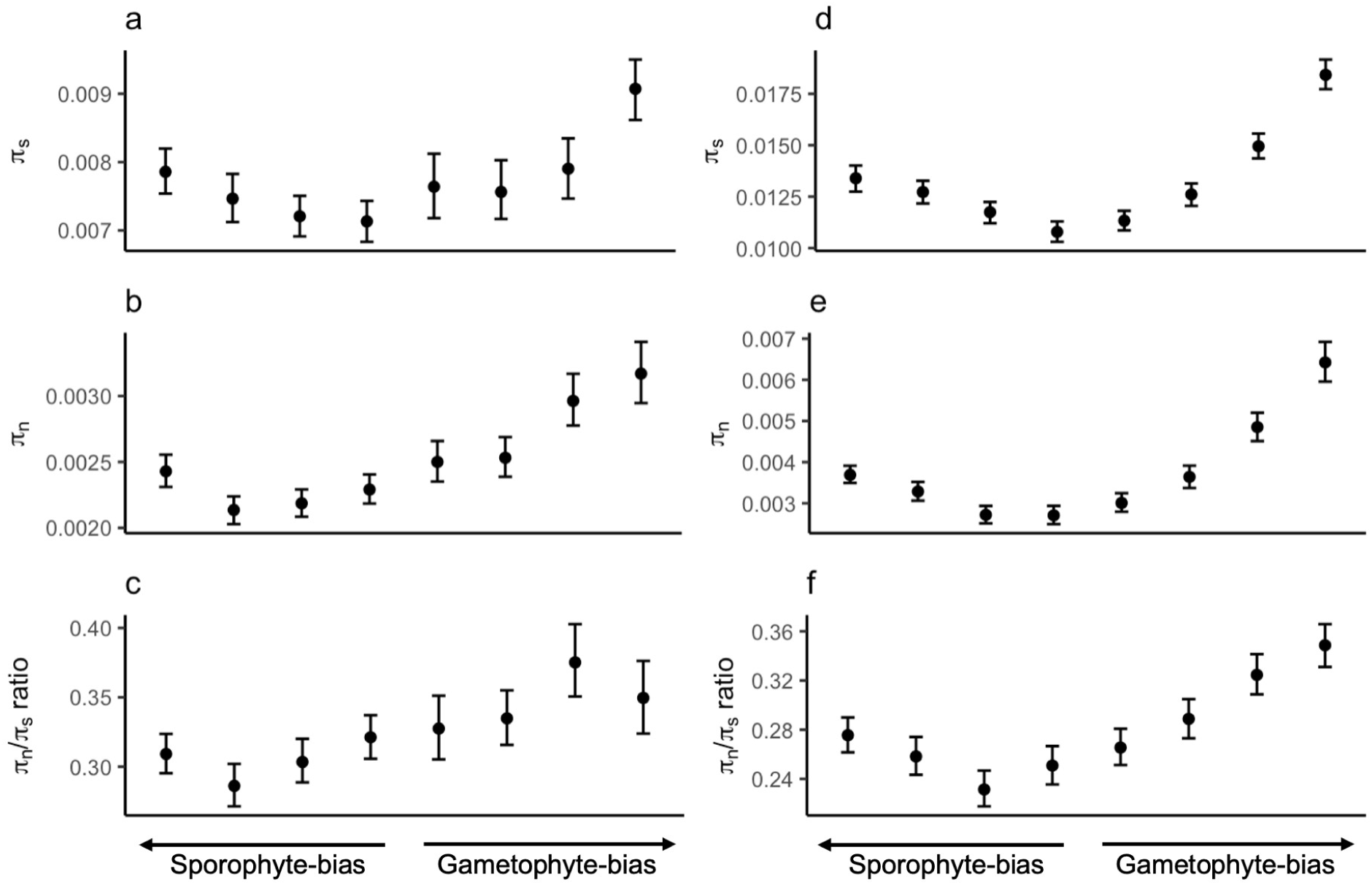
Effect of expression bias between life stages on nucleotide diversity in *R. hastatulus* (a-c) and *C. purpureus* (d-f). x-axis: quantiles of log2FoldChange between gametophyte and sporophyte expression. Number of genes in each bin: 1,009-1,847 for *R. hastatulus*; 1,943-2,564 for *C. purpureus*. The weighted mean and 95% confidence intervals are based on 1000 bootstraps of the original dataset. Note that the scales on the y-axis are different between species.

There are two possible explanations for the patterns of elevated *π*_s_ and *π*_n_ in the most gametophyte-biased genes, a greater prevalence of balancing selection under the assumption of neutrality of synonymous mutations or relaxed purifying selection on this set of genes. If there is relaxed purifying selection on gametophytic-biased genes, this could drive higher *π*_n_ and higher *π*_n_/*π*_s_. The observed *π*_n_/*π*_s_ ratios showed a similar trend as *π*_s_ and *π*_n_, especially in *C. purpureus*, with highly gametophyte-biased genes having the highest values (Figure 1c, 1f), suggesting either balancing selection, or a reduction in the strength of purifying selection on the most gametophyte-biased genes.

To further distinguish between the two alternative explanations for increased nucleotide diversity in gametophyte biased genes, we examined diversity patterns at genes specifically expressed in one of the two life stages, since these genes should experience the least pleiotropy across stages with reduced expression breadth. We compared *π*_s_, *π*_n_ and *π*_n_/*π*_s_ ratio for gametophyte-specific, sporophyte-specific and unbiased genes (Figure 2, S4; Table S8). The results are mostly consistent with Figure 1, with gametophyte-specific genes having a higher level of *π*_s_ and *π*_n_ than sporophyte-specific and unbiased genes in both species (Figure 2a, 2b, 2d, 2e), although the *π*_n_/*π*_s_ ratios were not elevated above unbiased genes in *R. hastatulus* as in *C. purpureus* (Figure 2c, 2f).

**Figure 2.**
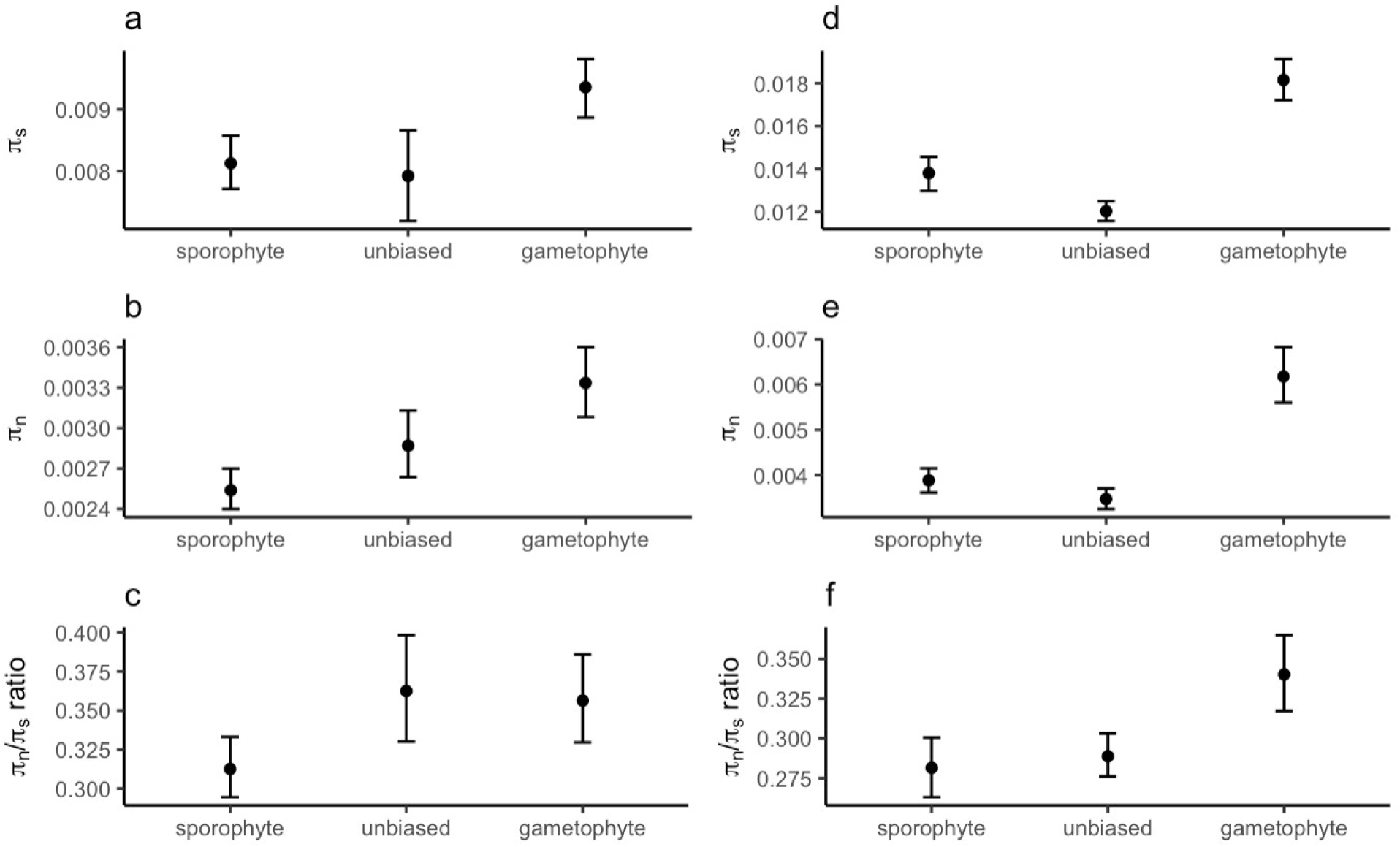
Weighted mean nucleotide diversity of gametophyte-specific, sporophyte-specific, unbiased genes in *R. hastatulus* (a-c) and *C. purpureus* (d-f). Number of genes in each category: 1149 (s), 920 (g), 524 (unbiased) for *R. hastatulus*; 1,232 (s), 1,217 (g), 3380 (unbiased) for *C. purpureus*. The weighted mean and 95% confidence intervals are based on 1000 bootstraps of the original dataset. Note that the scales on the y-axis are different between species.

One possible additional factor that could contribute to differences in the strength of purifying selection is if these genes differ in expression level. Gene expression has been shown to be a strong predictor of the strength of purifying selection genome-wide in many species (e.g., Urrutia and Hurst 2003; Slotte et al. 2011; Zhang and Yang 2015). We used this relation to further examine the possible effects of relaxed purifying selection on *π*_s_, and divided the genes into ten bins of expression level (baseMean across all samples) to examine how expression level affects *π*_s_, *π*_n_ and *π*_n_/*π*_s_ in gametophyte- and sporophyte-biased genes. As expression level increases, both *π*_n_ and *π*_s_ decrease, with a bigger decrease in *π*_n_ than *π*_s_ (Figure 3). Gametophyte-biased genes have higher *π*_s_, *π*_n_ and *π*_n_/*π*_s_ than sporophyte-biased genes, regardless of expression level in both species, except for the bins with overlapping confidence intervals in *π*_s_ of *R. hastatulus* (Figure 3). Our results suggest that expression level has a strong effect on patterns of diversity. Overall, contrary to our expectations, genes with overlapping expression levels did not show signals of elevated diversity due to balancing selection. Instead, heterogeneity in diversity likely reflects differences in the strength of purifying selection.

**Figure 3.**
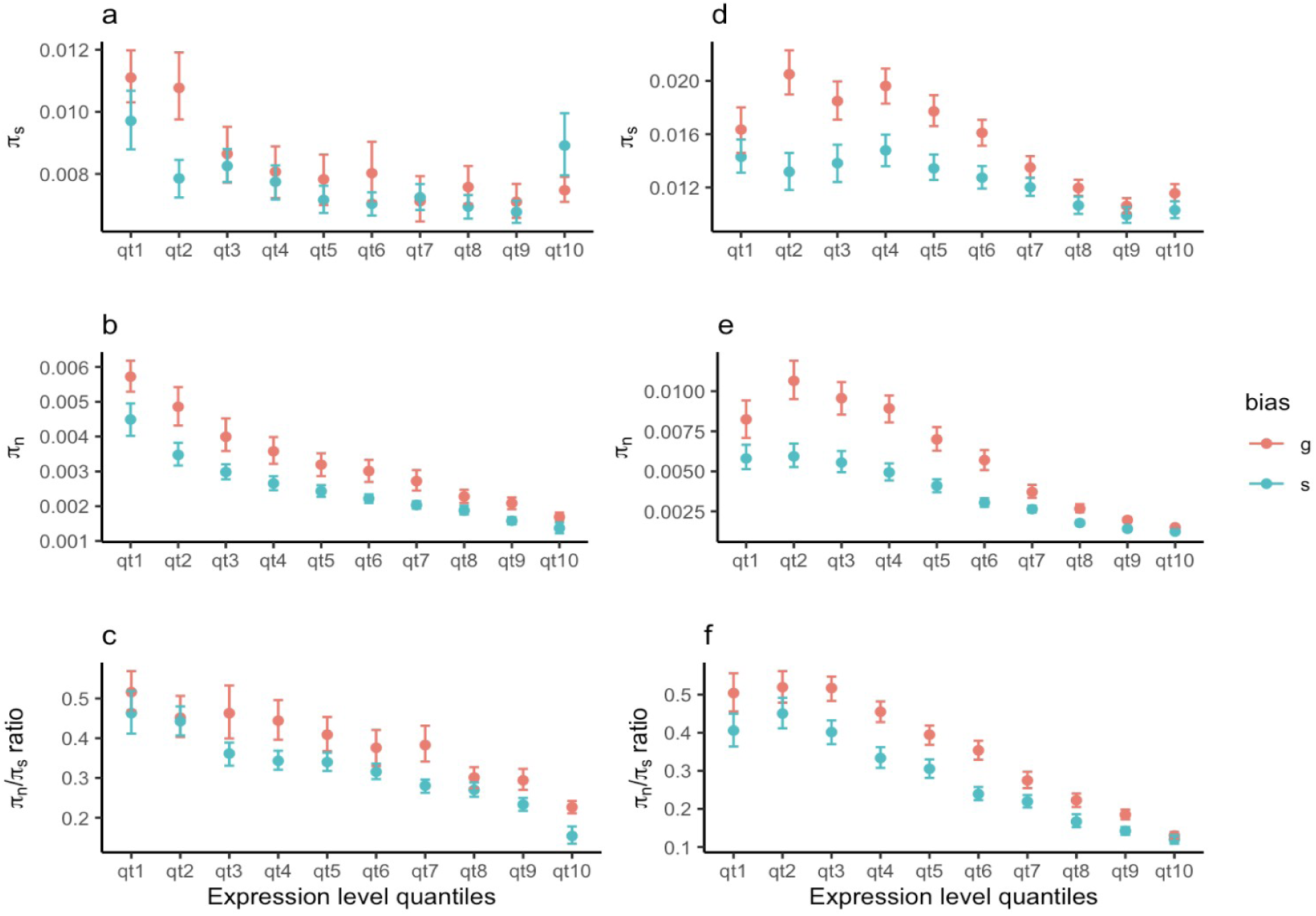
Effect of expression level on weighted mean nucleotide diversity in *R. hastatulus* (a-c) and *C. purpureus* (d-f). Number of genes in each bin: 686-1,464 for *R. hastatulus*; 1,034-2,328 for *C. purpureus*. The weighted mean and 95% confidence intervals are based on 500 bootstraps of the original dataset. qt1: lowest expression, qt10: highest expression. Note that the scales on the y-axis are different between species.

### Tajima’s D and expression bias between life stages

Tajima’s *D* is a metric summarizing information of the distribution of allele frequencies in a population (Tajima 1989) and is influenced by both selection and demographic history. Positive values are associated with balancing selection or population contraction and negative values indicate purifying selection or population expansion. We further tested the effect of expression bias between life stages on Tajima’s *D* at synonymous (*D*_s_) and nonsynonymous (*D*_n_) sites (Figure 4). In both species, *D*_n_ was smaller than *D*_s,_ consistent with purifying selection on nonsynonymous sites (Figure 4a v.s. 4b, 4c v.s. 4d). In *R. hastatulus*, both *D*_n_ and *D*_s_ were negative, *D*_s_ was higher in gametophyte-biased genes than unbiased and sporophyte-biased genes, as expected if they had a higher proportion of genes under balancing selection (Figure 4a). The negative values in *D*_n_ and *D*_s_ in *R. hastatulus* may possibly be attributed to population expansion, but we currently have no evidence on demographic history. In *C. purpureus*, the patterns of *D*_n_ were more similar to *π*_s_ and *π*_n_ than *D*_s,_ both *D*_s_ and *D*_n_ were higher in highly gametophyte-biased genes than unbiased genes (Figure 4c, 4d). The higher and more positive Tajima’s *D* values in *C. purpureus* compared to *R. hastatulus* likely resulted from differences in the two species’ demographic histories. We also compared *D*_s_ and *D*_n_ in gametophyte- and sporophyte-biased genes across expression level quantiles. In *R. hastatulus*, *D*_s_ and *D*_n_ remained roughly constant across expression level quantiles, whereas in *C. purpureus*, *D*_s_ and *D*_n_ decreased with higher expression (Figure S5). Gametophyte- and sporophyte-biased genes had similar levels of *D*_s_ and *D*_n_ across expression levels in both species. Combined with patterns of *π* (Figure 1-3) our results are consistent with the hypothesis that gametophyte-biased genes are under relaxed purifying selection.

**Figure 4.**
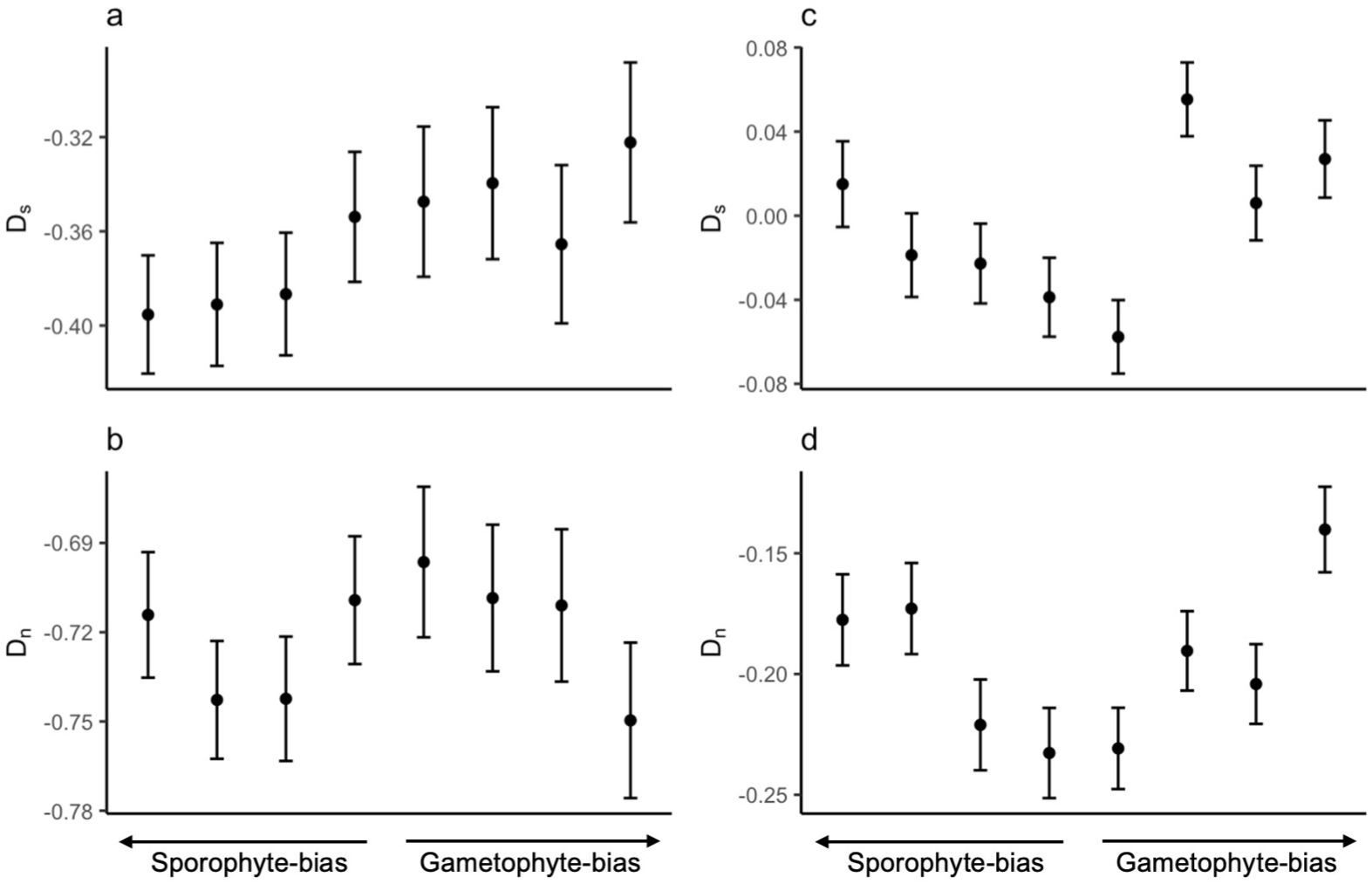
Effect of expression bias between life stages on Tajima’s *D* in *R. hastatulus* (a, b) and *C. purpureus* (c, d). x-axis: quantiles of log2FoldChange between gametophyte and sporophyte expression. Number of genes in each bin: 1,009-1,734 (*D*_s_), 1,197-1,966 (*D*_n_) for *R. hastatulus*; 2,138-2,534 (*D*_s_), 2,097-2,579 (*D*_n_) for *C. purpureus*. Error bars represent mean ± SEM across genes in each bin. Note that the scales on the y-axis are different between species.

### Distribution of fitness effects

Tajima’s *D* values reflect a summary of the site frequency spectrum rather than an explicit quantification of the strength of purifying selection on nonsynonymous sites. To address this, we estimated the distribution of fitness effect (DFE) to further understand the strength of purifying selection across life stages while controlling for expression level (Supplementary Methods). We first compared the DFE of gametophyte-specific, sporophyte-specific and unbiased genes (Figure S6, Table S9), then we binned the gene sets into four expression level quantiles (Figure S7, Table S10). In both species, higher expression level was associated with lower proportions of effectively neutral mutations (0 < *N_e_s* < 1), consistent with an increasing level of constraint due to higher expression level (Figure S7).

In *C. purpureus*, unbiased genes had a lower proportion of effectively neutral mutations and a higher proportion of strongly deleterious mutations than gametophyte- or sporophyte-specific genes. This finding is consistent with unbiased genes experiencing higher levels of constraint whereas stage-specific genes are more likely under relaxed purifying selection (Figure S6b). Similarly, in the highest expression level quantile (qt4), unbiased genes had a slightly lower proportion of effectively neutral mutations than gametophyte- or sporophyte-specific genes. However, in the other expression level quantiles the same pattern was not evident: unbiased genes had a higher proportion of effectively neutral mutations in the first quantile, and lower proportion of strongly deleterious mutations in the second quantile (Figure S7b). In *R. hastatulus*, gametophyte-specific and unbiased genes had similar proportions of effectively neutral and strongly deleterious mutations and sporophyte-specific genes had the lowest proportion of effectively neutral mutations and the highest proportion of strongly deleterious mutations (Figure S6a), consistent with Figure 2c. Across expression level quantiles, the confidence intervals mostly overlapped between gene sets and mutation categories in *R. hastatulus* (Figure S7a).

Based on the haploid purging hypothesis, we predicted that selection efficacy would be stronger in haploid than diploid phases. We therefore expected gametophyte-specific genes to have higher proportions of slightly (1 < *N_e_s* < 10) or strongly (*N_e_s* > 10) deleterious mutations, or a lower proportion of effectively neutral mutations (0 < *N_e_s* < 1). In contrast to previous studies (Arunkumar et al. 2013; Gossmann et al. 2014; Harrison et al. 2019; Gutiérrez-Valencia et al. 2021; Cervantes et al. 2023), our results in both species did not consistently support the haploid purging hypothesis. In *R. hastatulus*, gametophyte-specific genes had a higher proportion of strongly deleterious mutations than sporophyte-specific genes, while in *C. purpureus* the proportions were similar with overlapping confidence intervals (Figure S6). In the low to intermediate expression quantiles (qt1 to qt3), sporophyte-specific genes showed a slightly higher proportion of strongly deleterious mutations (*N_e_s* > 10) than gametophyte-specific genes in both species (Figure S7). In the highest expression quantile (qt4), gametophyte-specific genes had slightly higher proportions of strongly deleterious mutation (*N_e_s* > 10) than sporophyte-specific genes in both species, but the differences were not significant with overlapping confidence intervals (*R. hastatulus*: 67.19 - 70.21% in n, 65.38 - 68.66% in 2n; *C. purpureus*: 77.09 - 80.36% in n, 74.22 - 78.78 % in 2n, Figure S7, Table S10).

### Genome-wide scan for balancing selection

These overall patterns of diversity do not show a genome-wide signal of an enrichment of balancing selection on genes with overlapping expression between the life stages; however, it is possible that individual genes may still be subject to balancing selection. We therefore took an alternative approach by conducting a genome-wide scan for signals of balancing selection and explored whether they were enriched for genes that might be subject to antagonistic pleiotropy between life stages. We applied a composite likelihood ratio test to scan the genome for candidate genes under balancing selection based on *B*_2_ statistics that account for both increased polymorphism density and an excess of intermediate-frequency alleles (Cheng and DeGiorgio 2019, 2020, 2022). In *R. hastatulus,* a total number of 4,945,561 sites were tested in windows based on genetic positions (Figure 5). We found 79,431 sites with strong signals of balancing selection across all chromosomes (CLR > 9.5), located at 449 genes, 405 of which had evidence of expression. The candidate genes were only located in regions with considerable recombination where genetic positions increased rapidly with physical positions, based on our genetic maps (Figure 5, Figure S11). However, among the genes found in regions with evidence of balancing selection, 138 were located within inversion polymorphisms identified from two phased whole genome assemblies of *R. hastatulus* on chromosomes A2 (63), A3 (74), and A4 (1) (Sacchi et al. 2024; Sacchi et al, in prep), consisting of more than 30% of all candidate genes; whereas these inversions contained 16.7% of genes across the genome. Because inversion heterozygotes are likely to experience reduced recombination, these candidate genes may be subject to non-independent signals of balanced polymorphism. In *C. purpureus*, the scans were performed in physical position windows as site-specific recombination rates were unavailable (Figure S8). We tested 7,743,902 sites and identified 339,375 sites under balancing selection (CLR > 956.2), located at 963 genes, 807 of which had evidence of expression in this study. In *R. hastatulus*, as expected candidate genes under balancing selection had significantly higher *π*_s_, and significantly lower *π*_n_/*π*_s_ than genes that were not under balancing selection (Welch Two Sample *t*-test, *p*-value = 0.0001511, 0.0003204, respectively). In *C. purpureus*, there was no significant differences in *π*_s_ or *π*_n_/*π*_s_ between genes under balancing selection and gene that were not (Welch Two Sample *t*-test, *p*-value = 0.6029; Two Sample *t*-test, *p*-value = 0.5956, respectively). We compared the site frequency spectrum (SFS) of all sites tested and the sites under balancing selection (Figure S9, S10) and found sites under balancing selection had significantly higher minor allele frequencies than the input data in both species (Mann-Whitney *U* test, *p*-value < 2.2e-16). Note that this is expected, given the use of the site frequency spectrum in the model-based inference. The differences in the shape of SFS in *C. purpureus* is more prominent with a higher proportion of common alleles than in *R. hastatulus* (Figure S9, S10).

**Figure 5.**
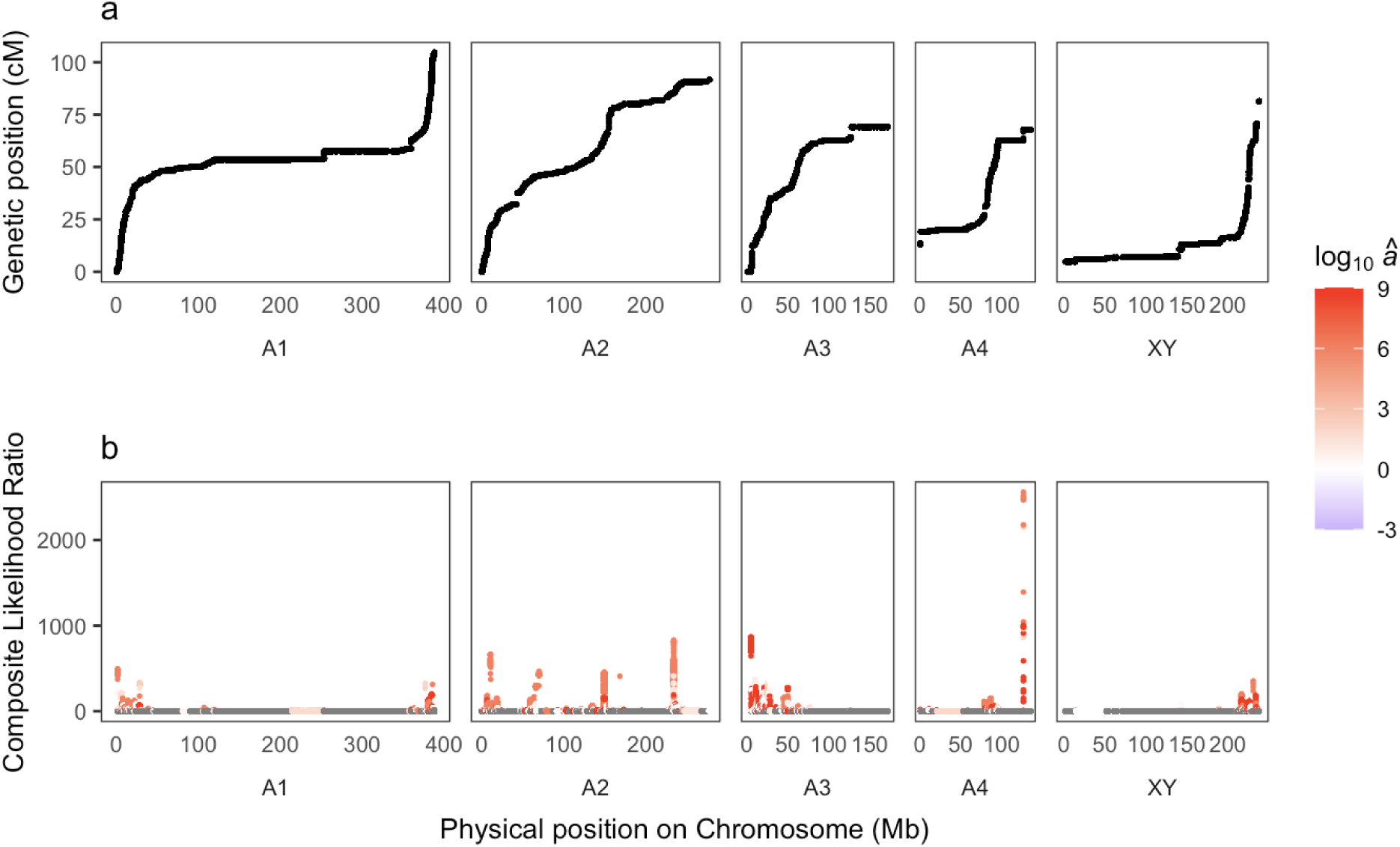
Genome-wide scan of balancing selection in *R. hastatulus*. y-axis: genetic positions of sites being tested (a), composite likelihood ratio (b). *â*: estimated dispersion parameter, a positive log_10_*â* value suggests balancing selection.

We performed contingency tests on the number of genes in different expression bias categories (gametophyte-biased, sporophyte-biased, unbiased) for genes under balancing selection and across the whole genome (Tables S11). In both species, we found no enrichment of any category of expression bias under long-term balancing selection compared to the whole genome (Fisher’s exact test, *p* > 0.05). We examined the gene functions for all gametophyte-biased genes in regions under balancing selection (Table S12, S13). We performed GO enrichment after excluding the genes found within inversion polymorphisms in *R. hastatulus*, since the suppression of recombination between inversion types can influence our ability to find functional enrichments of causal loci. The GO enrichment results after excluding inversions were not compelling (Table S12). In *R. hastatulus*, two genes under balancing selection had probable disease resistance functions based on orthologs to *Arabidopsis thaliana* (At4g19060, At4g14610). In *C. purpureus*, two genes under balancing selection were expressed in both life stages with high gametophyte bias and have sexual reproduction functions (log2FoldChange > 3, GO:0019953). These genes were also in the top 10% quantile of *π*_s_.

## Discussion

We combined population genomic and gene expression analyses to test for balancing selection due to conflict between life stages in *R. hastatulus* and *C. purpureus*. If balancing selection is primarily driven by antagonistic pleiotropy between life stages, we would expect stronger signals of balancing selection in genes expressed in both life stages. Instead, we found elevated *π*_s_, *π*_n_ and *D*_n_ in highly gametophyte-biased genes in both species. Combined with the observations that *π*_s_, *π*_n_ and *π*_n_/*π*_s_ all decreased with increased expression levels, the elevated diversity suggests relaxed purifying selection rather than balancing selection. DFE analyses did not show stronger purifying selection in gametophyte-than sporophyte-specific genes. Lastly, genome-wide scans identified hundreds of candidate genes under balancing selection, but these genes were not enriched for gametophyte-biased expression. Below we discuss how these findings inform our understanding of plant gametophytic selection and the challenges of alternative approaches for detecting balancing selection.

### Widespread relaxed purifying selection

Balancing selection driven by sexual antagonism has been tested using genetic diversity and sex-biased expression (Cheng and Kirkpatrick 2016; Kasimatis et al. 2017; Mank 2017; Wright et al. 2018; Sayadi et al. 2019), showing elevated diversity in weakly sex-biased genes consistent with ongoing conflict. We predicted similar patterns for life stages, however, we found higher *π*_s_, *π*_n_ and *D*_n_ in highly gametophyte-biased genes (Figure 1), which could result from balancing selection or relaxed purifying selection. As highly expressed genes are under strong selective constraint (e.g., Urrutia and Hurst 2003; Slotte et al. 2011; Zhang and Yang 2015), high *π*_s_ in gametophyte-biased genes with high expression level would suggest balancing selection. Since *π*_s_ decreased with higher expression level (Figure 3), the elevated *π*_s_ in gametophyte-biased genes suggests relaxed constraint on both nonsynonymous and synonymous sites (Zhang and Qian 2025) is the more plausible explanation rather than widespread intralocus conflict.

Higher *π*_s_ under relaxed purifying selection could stem from weaker background selection and/or less direct selection on synonymous sites themselves. Background selection reduces neutral nucleotide diversity due to linkage to deleterious mutations, with greater reductions in more constrained regions (Elyashiv et al. 2016). Highly gametophyte-biased genes may be under weaker background selection, elevating *π*_s_ and *π*_n_. Though synonymous mutations are generally assumed to be neutral (King and Jukes 1969), studies in fruit flies and yeast showed many synonymous mutations can be deleterious (Lawrie et al. 2013; Shen et al. 2022; Zhang and Qian 2025; but see Kruglyak et al. 2023), likely due to codon usage bias (Plotkin and Kudla 2011; Hunt et al. 2014; Zhou et al. 2016). In *Marchantia polymorpha*, relaxed purifying selection and lower codon usage bias were found in sporophyte-biased genes (Shen et al. 2024). In our case, both synonymous and nonsynonymous sites may be constrained but less so in highly gametophyte-biased genes, resulting in elevated *π*_s_ and *π*_n_.

Expression level and breadth may have a stronger effect on selection efficacy than antagonistic pleiotropy across life stages. Our results showed higher expression level decreased diversity and the proportions of effectively neutral mutations (Figure 3, Figure S7). Gametophyte- and sporophyte-biased genes likely have more specialized functions and reduced expression breadth than unbiased genes, particularly pollen-biased genes (Honys and Twell 2003), which could explain their relaxed constraint. However, limited tissue sampling in *R. hastatulus* (pollen and leaf from a specific developmental stage) restricted our assessment of expression breadth and may have affected functional enrichment (Table S5). Nonetheless, studies in *Arabidopsis thaliana* have indicated that pollen expression is highly distinct from that of sporophytic tissues, suggesting that pollen-biased genes are likely strongly enriched for gametophytic functions (Rutley and Twell 2015). The lack of female gametophytic tissue (i.e., ovules) in *R. hastatulus* limits our ability to detect potential sexual conflict within the gametophytic stage and between sporophytes and female gametophytes. Additionally, sperm competition can contribute to the maintenance of genetic variation (Clark 2002; Dapper and Wade 2016). Sperm and vegetative cells in pollen might experience different selection pressures, which could explain the patterns of DFE in *R. hastatulus* (Arunkumar et al. 2013; Gossmann et al. 2014; Gutiérrez-Valencia et al. 2022). In contrast, *C. purpureus* expression data included many cell types, the more symmetric expression bias-diversity relationship (Figure 1) may reflect smaller differences in expression breadth between gametophytes and sporophytes. Future studies should benefit from a wider and more complete sampling of tissues from different plant species, e.g., including roots in angiosperms like *R. hastatulus*.

Despite different sampling strategies, genome properties, and contrasting predominance of the gametophytic stage, both species showed elevated diversity in highly gametophyte-biased genes. The whole-genome resequencing data of *C. purpureus* came from geographically diverse isolates, while the whole-genome sequencing data of *R. hastatulus* were from a single population (see Material and Methods). Although unlikely to drive a genome-wide pattern, spatially variable selection in *C. purpureus* may maintain different locally adapted alleles especially in the long-lived gametophytes, increasing *π* and Tajima’s *D*. The large non-recombining regions in *R. hastatulus* affects gene density and thus patterns of genetic diversity (Rifkin et al. 2022). In contrast, bryophytes like *C. purpureus* show higher recombination rates and more uniform gene density across the genome (Gaut et al. 2007; Bowman et al. 2017; Lang et al. 2018; Healey et al. 2023). The consistent patterns across species support relaxed purifying selection as the most plausible explanation. In contrast to antagonistic pleiotropy, the prevalence of synergistic pleiotropy between life stages may explain the stronger signals of purifying selection on unbiased genes, i.e., deleterious mutations in these genes are selected against in both life stages. This concordant selection could dominate genome-wide patterns of diversity, and is consistent with breeding studies in which selection on pollen can have synergistic pleiotropic effects on sporophytic fitness (Hormaza and Herrero 1992).

### Genome-wide scan for balancing selection

The maintenance of genetic variation through balancing selection is a central to evolutionary biology, yet detecting balancing selection genome-wide remains challenging (Ruzicka et al. 2025). Traditional methods rely on the HKA test (Hudson et al. 1987) and summary statistics like Tajima’s *D*, which may lack power compared to composite likelihood ratio (CLR) based test (Bitarello et al. 2023). We used the CLR-based *B* statistics, which adjusts window size based on the data and can integrate multiple signatures including an excess of common alleles and increased density of polymorphisms (Cheng and DeGiorgio 2019, 2020, 2022). However, even with improved performance, *B* statistics have limited power when used to detect a single signature of selection and determining appropriate window sizes is challenging without a genetic map. Genome-wide assessments of selection are affected by confounding factors such as demographic history and variation in recombination rates. Limited knowledge of the functional annotation in the study system further complicates the interpretation of candidate sites under selection. Detection power depends on the underlying selection model; for example, Flintham et al. (2025) showed that balancing selection driven by sexual conflict is difficult to detect in genomics, as its polymorphism signature is transient under polygenic selection. Similar challenges are expected when detecting balancing selection due to intralocus conflict between life stages.

Although we did not find a genome-wide signal of intralocus conflict between life stages, we performed functional enrichment of candidate genes under balancing selection to test if they are subject to conflict. We identified 499 and 964 genes under balancing selection in *R. hastatulus* and *C. purpureus*, respectively. Similar approaches should be informative to test whether sexually antagonistic SNPs are under balancing selection (Ruzicka et al. 2019). Our results show over 30% of candidate genes for balancing selection (138 genes) in *R. hastatulus* are located within inversion polymorphisms, consistent with findings from humans (Giner-Delgado et al. 2019). Since recombination is suppressed in inversion heterozygotes, many of these 138 genes may show signals of balancing selection due to linkage disequilibrium rather than being direct targets of selection. This linkage effect makes it difficult to distinguish focal genes under balancing selection from nearby unselected genes, which potentially affects our functional enrichment results (Table S12, S13). Similarly, we did not observe an enrichment of gametophyte-biased genes among those under balancing selection. However, in *C. purpureus*, we identified two gametophyte-biased genes that show signals of balancing selection; studies of such genes could shed light on the nature of any intergenerational antagonistic pleiotropy.

### Gene expression profiles in species with biphasic life cycles

The differentially expressed genes between life stages and their functional enrichment suggest distinct ecological and reproductive roles for gametophytes and sporophytes. In *R. hastatulus*, the smaller overlap and greater differentiation in gene expression between life stages (Table 1) may reflect the evolutionary reduction of the gametophytic stage in angiosperms, where gametophytes become short-lived, highly specialized, and structurally reduced compared to those in bryophytes (Szövényi et al. 2011). However, differences in RNA sequencing technologies, sample sizes and gene annotations methods between studies limit direct comparisons. Broader comparative analyses across major land plant lineages, such as gymnosperms (Cervantes 2023), using standardized methods would help clarify how life stage complexity and duration affect the extent of overlapping and differential gene expression.

## Methods

### Plant materials

For *R. hastatulus*, we collected seeds from a population in Rosebud, Texas, US (Pickup and Barrett 2013) and planted seedlings in a glasshouse at Earth Science Center, University of Toronto. We collected pollen grains from the male F_1_ plants for RNA isolation based on a modified protocol from Lu (2011). We collected male leaf tissues for RNA isolation and female leaf tissues for DNA isolation using F_1_ plants, see supplementary methods for details. We used the standard protocols of Qiagen Plant Mini kit and Sigma Aldrich Spectrum Plant Total RNA Kits for DNA and RNA isolation, respectively. All libraries were prepared and sequenced at the Genome Quebec Innovation Centre in Montréal, Canada. The DNA samples were sequenced at a depth of 10-15X. The median read numbers for DNA and RNA sequences are 72,907,022 (range: 64,585,743 - 88,888,320) and 23,412,838 (range: 18,618,763 - 58,151,564), respectively (Table S1, S2). Our final sample sizes were 20 female leaf DNA, 77 male leaf RNA, and 75 pollen RNA.

For *C. purpureus*, gametophyte DNA sequences including 18 samples at 6 locations are available from Carey et al. (2021c). All RNA samples were taken from juvenile (protonema) and mature (gametophore) tissues at the gametophyte stage. Gametophyte RNA sequences from 3 male-female sibling pairs at 2 developmental stages (each with 3 biological replicates) are available from Carey et al. (2021a). We generated sporophyte tissues for RNA sequencing using the same methods and at the same time with Carey et al. (2021a). The sporophyte RNA sequences included 26 samples from 6 crosses, the number of replicates per cross is between 2 and 10 (Table S3). We removed one sporophyte RNA sample from our analyses because it had low coverage. The median read number for the sporophyte RNA sequences is 52,038,814 (range: 33,030,143 - 64,278,575) (Table S3).

### RNA Seq data and expression analysis

For *R. hastatulus*, we used the genome assembly and genome annotation from Rifkin et al. (2022) and assessed the quality of RNA reads using FastQC (Andrews 2010). In *C. purpureus*, we used a hybrid genome consisting of the male R40 genome assembly (autosomes, the V sex chromosome) and the U sex chromosome from the female GG1 genome assembly (Carey et al. 2021a). We trimmed adaptor sequences and filtered out low-quality reads using trimmomatic for all RNA samples (Bolger et al. 2014), and trimmed polyG sequences using fastp for the sporophyte RNA samples (Chen 2023). We used FastQC to assess the quality of RNA reads before and after filtering (Andrews 2010). In *C. purpureus*, we excluded four sporophyte samples for being an outlier in PCA from the majority of sporophyte samples; our final sample size for sporophyte RNA samples is 21.

In both species, we mapped the RNA reads to the genome assembly using STAR in two pass mode (Dobin et al. 2013, v2.7.6a). We sorted the alignment SAM files and added read groups using PicardTools (http://broadinstitute.github.io/picard/). We generated raw read counts for each gene using featureCount (Liao et al. 2014) and then normalized the raw read counts by library depth using DESeq2 (Love et al. 2014). In the expression analysis of *C. purpureus*, we included both male and female gametophyte samples for autosomal genes and only included female or male gametophyte samples for U and V genes, respectively. A gene was identified as expressed if the mean normalized read count for either life stages was > 5; we only kept expressed genes in the subsequent analyses. We calculated expression ratios between life stages for each gene by dividing the mean normalized read counts at each life stage. The criteria for differentially expressed genes were Benjamini-Hochberg adjusted *p*-value < 0.1, corresponding to a false discovery rate of 0.1 and FC>2 (Benjamini and Hochberg 1995), allowing for a sensitive characterization of biological signals and a reasonable control for false positives. A gene was considered tissue-specific if: a) it was in the 10% or 90% quantiles of expression ratio among all significantly differentially expressed genes (adjusted *p*-value < 0.1), and b) it had an absolute value of log2FoldChange > 2. We performed GO enrichment for biased and specific genes using topGO, with expressed genes from each species serving as the gene universe for comparison. (Alexa and Rahnenfuhrer 2023). We divided the genes into four bins based on quantiles of log2FoldChange for gametophyte-biased (log2FoldChange > 0) and sporophyte-biased (log2FoldChange > 0) genes separately. We defined unbiased genes as the middle two bins with lowest absolute value of log2FoldChange as well as an adjusted *p*-value ≥ 0.1. We divided the expressed genes into four or ten quantiles of expression levels and used the same cutoff for later analyses.

### DNA Seq data and variant calling

For *R. hastatulus*, we assessed the quality of DNA raw reads using FastQC (Andrews 2010) and mapped the reads to the genome assembly using bwa-mem2 (Vasimuddin et al. 2019). We sorted the BAM files using SAMtools (Danecek et al. 2021), added read groups, and removed PCR duplicates using PicardTools (http://broadinstitute.github.io/picard/). We called SNPs jointly on all samples using freebayes (Garrison and Marth 2012) with invariant sites saved in the output (-- report-monomorphic). The variant and invariant sites were filtered separately using VCFtools (Danecek et al. 2011), and concatenated together using BCFtools (Danecek et al. 2021). For both variant and invariant sites we removed sites with a proportion of missing data > 0.8 or with a mean read depth per site <10 or >40. For variant sites, we only kept bi-allelic SNPs with genotype quality > 30.

For *C. purpureus*, we removed adaptor sequences using trimmomatic (LEADING:3 TRAILING:3 SLIDINGWINDOW:10:30 MINLEN:40) (Bolger et al. 2014), and performed quality control using FastQC before and after trimming (Andrews 2010). We masked the V sex chromosome when mapping female samples and masked the U sex chromosome when mapping male samples using BEDTools maskfasta (Quinlan and Hall 2010). The read mapping and BAM file processing methods are the same as described above in *R. hastatulus*. We called SNPs on all samples jointly using BCFtools mpileup with a ploidy of 1 and the -B option (Danecek et al. 2021). We only kept the VCF for autosomal genes for the subsequent analysis, as the U and V sex chromosomes have low recombination (Carey et al. 2021a, 2021c). We performed VCF filtering using BCFtools (Danecek et al. 2021). We removed sites with a combined read depth < 5 and filtered variant and invariant sites separately. For variant sites, we kept bi-allelic SNPs and removed sites with low quality or mapping score (QUAL<30 && MQ<30). Finally, we removed sites with a proportion of missing data > 0.8 for both invariant and variant sites.

### Diversity statistics

In both species we identified 0-fold and 4-fold degenerate sites in the genome using the script codingSiteTypes.py (https://github.com/simonhmartin/genomics_general/blob/master/codingSiteTypes.py, accessed in 2020). We intersected the list of 0-fold and 4-fold sites with the filtered VCFs to generate 0-fold and 4-fold VCFs for population genetic analyses. The numbers of sites (variant and invariant) in each VCF are 11,432,804 (0-fold) and 2,891,775 (4-fold) in *R. hastatulus*; 17,394,329 (0-fold) and 4,759,819 (4-fold) in *C. purpureus*. We used pixy to calculate the 0-fold and 4-fold average per site nucleotide diversity for each gene across the genome (Korunes and Samuk 2021, v1.2.6.beta1). Only genes with a minimum of 50 sites for both 4-fold and 0-fold nucleotide diversity were kept in the subsequent analyses.

We used the script popgenWindows.py (https://github.com/simonhmartin/genomics_general/blob/master/popgenWindows.py, accessed in 2020) to calculate Tajima’s *D* for each gene based on the variant-only VCFs. We calculated the weighted average nucleotide diversity for each bin based on the number of sites for each gene (see supplementary methods). We bootstrapped the genes in each bin 1000 times to generate 95% confidence intervals. We calculated the weighted average nucleotide diversity (mean and 95% CI) in the same way for gametophyte-specific, sporophyte-specific and unbiased genes. We removed zeros for *π*_s_ and *π*_n_ to calculate the *π*_n_/*π*_s_ ratio for each gene.

All DNA samples in *R. hastatulus* are female, this reduces the bias in nucleotide diversity estimates due to the divergence between the X and Y sex chromosomes in males. In *C. purpureus*, nucleotide diversity was estimated from 16 DNA samples (8 males and 8 females) from 5 locations, see sample information from (2021a).

### Balancing selection scan

We used BalLeRMix+ with default settings to perform genomic scans for balancing selection based on *B*_2_ statistics, incorporating derived allele frequency, polymorphism and divergence information (Cheng and DeGiorgio 2019, 2020, 2022). We used an outgroup to infer the ancestral state for both species (supplementary methods). A high CLR indicates strong signal for balancing selection. In *R. hastatulus*, the scans were performed on each chromosome using the genetic positions. In *C. purpureus*, the scans were performed in 1 Mb sliding windows based on the physical positions on each chromosome, using a default recombination rate of 1 cM/Mb. The threshold for signature of balancing selection was set as the 5% quantile in the non-zero values of composite likelihood ratio. We calculated the maximum CLR per gene. We identified the genes that had at least one site with strong signal of balancing selection and also with evidence of expression in this study. We performed GO enrichment for candidate balancing selection genes using topGO, with expressed genes as the gene universe for comparison (Alexa and Rahnenfuhrer 2023). For candidate balancing selection genes with gametophyte bias, we used gametophyte-biased genes as the comparison to test the functional enrichment for balancing selection.

## Data availability

Raw reads of *R. hastatulus* DNA and RNA Seq are available in NCBI SRA database under BioProject PRJNA744278 (to be released on Aug 1, 2025 or upon publication). Raw reads of *C. purpureus* sporophyte RNA Seq are available through JGI genome portal, see project IDs in supplementary Table S3. Scripts used in this study are available at https://github.com/imengyuan/lifestage_conflict.

## Acknowledgements

We thank Bill Cole and Thomas Gludovacz for help with plant maintenance, Andrew Clemens for assistance with *C. purpureus* sporophyte cultivation, and Tyler Kent for help with interpolating recombination rate for *R. hastatulus*. This work was supported by NSERC Discovery Grants awarded to JRS, SIW and SCHB, and NSF and NASA awards to SFM. The *C. purpureus* sequencing (proposal: 10.46936/10.25585/60007219) conducted by the U.S. Department of Energy Joint Genome Institute (https://ror.org/04xm1d337), a DOE Office of Science User Facility, was supported by the Office of Science of the U.S. Department of Energy operated under Contract No. DE-AC02-05CH11231. We thank Vincent Castric and two anonymous reviewers for comments on the manuscript.

