## Supplemental methods and figures for "Testing for the genomic footprint of conflict between life stages in an angiosperm and moss species"

### Supplementary methods

*Plant materials for Rumex hastatulus*

After flowering, we paired one female and one male individual randomly and moved them to a mini plastic chamber for crossing. We collected mature seeds from the crosses to generate $F_1$ plants. We collected leaf and pollen tissues from 77 male $F_1$ plants for RNA isolation and leaf tissues from 20 female $F_1$ plants for DNA isolation. Growth conditions and leaf tissue collection methods are available in Rifkin et al. (2022). We collected 2-3 mature inflorescences per male individual into a 15 mL falcon tube and added ice-cold 0.5 M mannitol solution till the tube was full. We vortexed the tube vigorously for 1 minute to release the pollen grains, filtered the pollen suspension through a 30 µm nylon mesh, and centrifuged the tube at 450 x g for 5 minutes (4 °C). We repeated the above step with fresh mannitol solution. We transferred the final pollen pellets to a 2 mL centrifuge tube that was flash frozen in liquid nitrogen and stored under -80 °C.

*Weighting in nucleotide diversity*

We multiplied the per-site average of $\pi_s$ by the number of synonymous sites for each gene and summed the resulting $\pi_s$ and the number of synonymous sites across all genes of the gene set, we then divided the sum of $\pi_s$ and the sum of number of synonymous sites to get the weighted average $\pi_s$ for the gene set. We repeated the same steps for nonsynonymous sites to get the weighted average of $\pi_n$, and $\pi_n/\pi_s$ ratio of each gene set. We resampled the genes 1000 times with replacement within each gene set to generate the mean and 95% confidence intervals for the weighted average $\pi_s$, $\pi_n$ and $\pi_n/\pi_s$. We also calculated the mean and SEM of per-site average of $\pi_s$ and $\pi_n$ across genes for each gene set to examine the effect of weighting.

*Distribution of fitness effect (DFE) analysis*

We removed sites with missing data from the 0-fold and 4-fold VCFs used in the diversity analysis. The number of sites in different VCFs reduced to 11,046,527 (0-fold) and 2,796,108 (4-fold) in *R. hastatulus*; 16,453,490 (0-fold) and 4,503,087 (4-fold) in *C. purpureus*. At each site, *R. hastatulus* has 40 alleles from 20 female sporophytes and *C. purpureus* has 16 alleles from 8 female and 8 male gametophytes. We used the 0-fold and 4-fold VCFs for different gene sets (e.g., gametophyte- and sporophyte-specific genes) to generate the 0-fold and 4-fold site frequency spectra. We estimated the distribution of fitness effects of new mutations for different gene sets using the DFE-alpha program with default setting (Keightley and Eyre-Walker 2007). Mean and 95% confidence intervals were generated by bootstrapping the sites in different gene sets 200 times.

*Input file for balancing selection scan*

We performed whole genome alignment between *R. hastatulus* and the close relative *R. bucephalophorus* using anchorwave (Song et al. 2022). The ancestral state of each site is determined by the allele in *R. bucephalophorus*, we calculated the derived allele frequency at each site in *R. hastatulus* accordingly. We interpolated the recombination rate for each site using the recombination maps from Rifkin et al. (2022). The interpolation was conducted using either a splinefun function in the R package stats, or a linear function if the sites are out of the range of the input recombination map (https://github.com/tvkent/BioinformaticsUtils/blob/master/interpolate.R) (Kent 2023). The derived allele frequencies in *C. purpureus* were calculated using Chile individuals (1 female, 1 male) as the outgroup (Carey et al. 2021).

*Inversions in R. hastatulus*

Using phased genome assemblies from the XYY cytotype (Sacchi et al. 2024), and the XY cytotype of *R. hastatulus* (Humphries et al, *in prep*) we used comparative pairwise dotplots based on syntenic gene positions in COGE (Lyons and Freeling 2008) to identify large (>1MB) inversion polymorphisms between haplotypes on the autosomes. Given evidence for shared inversion polymorphisms between the cytotype populations (Sacchi et al. 2024), we recorded the locations of all inverted regions, including those that are heterozygous between haplotypes within the XYY cytotype. These regional positions were localized on the reference genome used in this assembly using COGE (Lyons and Freeling 2008) for analysis of enrichment of balancing selection signals.

**Supplementary tables**

Table S1. *Rumex hastatulus* leaf and pollen RNA sequencing data.

Table S2. *Rumex hastatulus* leaf DNA sequencing data.

Table S3. *Ceratodon purpureus* sporophyte RNA sequencing data.

Table S4. Number of life-stage biased or specific genes in *Rumex hastatulus* and *Ceratodon purpureus.*

Table S5. GO enrichment for biased and specific genes between life stages in *Rumex hastatulus.*

Table S6. GO enrichment for biased and specific genes between life stages in *Ceratodon purpureus.*

Table S7. Estimates of nucleotide diversity in bins of expression bias in *Rumex hastatulus* and *Ceratodon purpureus.*

Table S8. Estimates of nucleotide diversity in different gene sets in *Rumex hastatulus* and *Ceratodon purpureus.*

Table S9. DFE estimates of life stage-specific genes in *Rumex hastatulus* and *Ceratodon purpureus.*

Table S10. DFE estimates of life stage-specific genes across expression level quantiles in *Rumex hastatulus* and *Ceratodon purpureus.*

Table S11. Number of biased genes under balancing selection vs across the whole genome.

Table S12. GO enrichment for genes under balancing selection in *Rumex hastatulus.*

Table S13. GO enrichment for genes under balancing selection in *Ceratodon purpureus.*

**Supplementary figures**

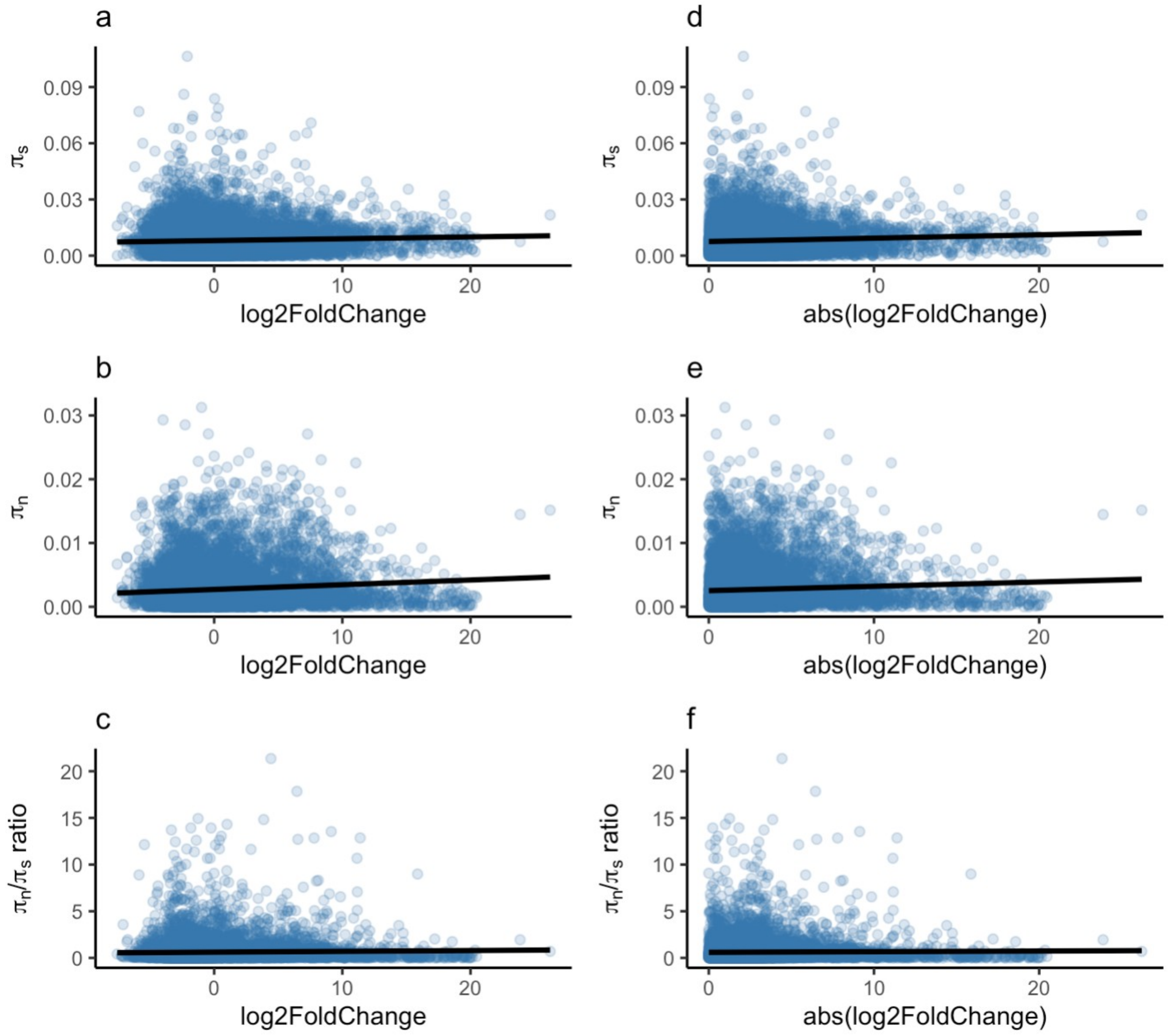


Figure S1. Linear regression between expression bias and nucleotide diversity in *R. hastatulus*. x-axis: log2FoldChange (a-c) or the absolute value of log2FoldChange (d-f). We removed zeros for $\pi_s$ and $\pi_n$ to calculate the $\pi_n/\pi_s$ ratio for each gene. The slopes and p-values for $\pi$ are all < 0.001 and < 10-8, respectively. The slopes for the $\pi_n/\pi_s$ ratio are both < 0.01, p-value = 0.00133 (c), 0.0766 (f).

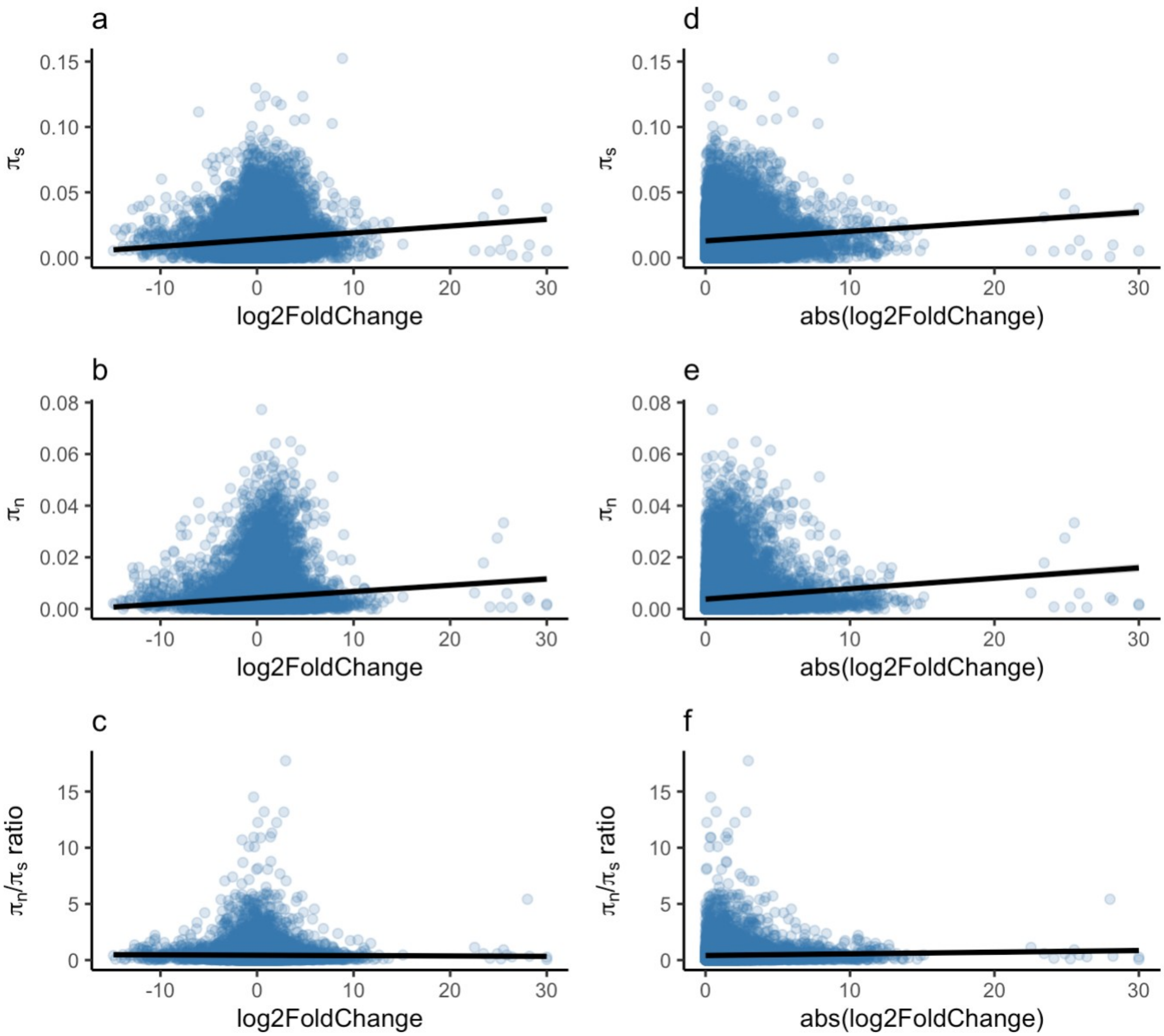


Figure S2. Linear regression between expression bias and nucleotide diversity in *C. purpureus*. x-axis: log2FoldChange (a-c) or the absolute value of log2FoldChange (d-f). We removed zeros for $\pi_s$ and $\pi_n$ to calculate the $\pi_n/\pi_s$ ratio for each gene. The slopes and p-values for $\pi$ are all < 0.001 and < 10-16, respectively. For the $\pi_n/\pi_s$ ratio: slope = -0.003242, p-value = 0.176 (c); slope = 0.014604, p-value = 1.73e-06 (f).

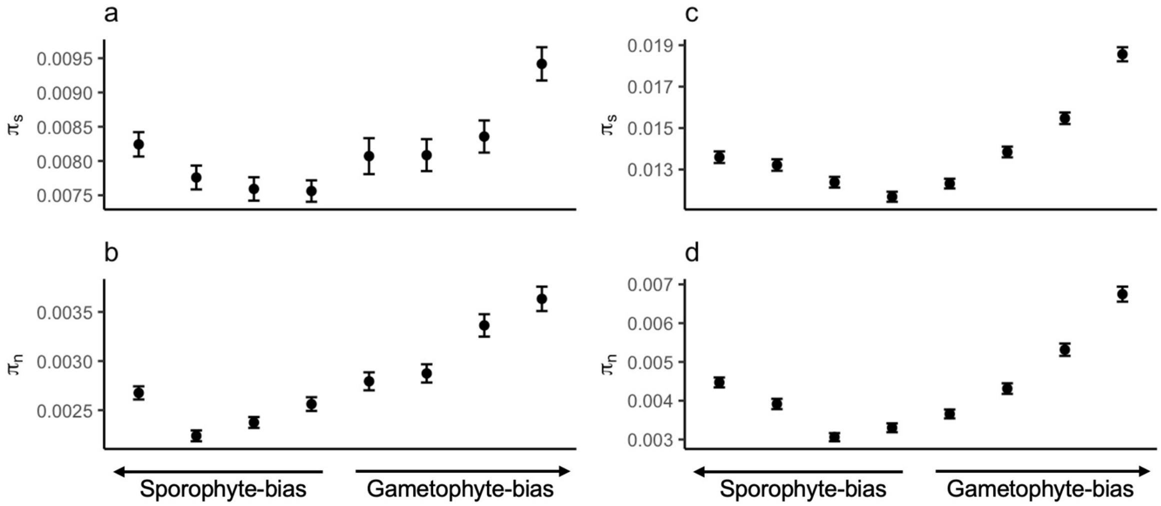


Figure S3. Effect of expression bias between life stages on nucleotide diversity in *R. hastatulus* (a-b) and *C. purpureus* (c-d). The numbers of genes in each bin are the same as Figure 1. Error bars represent mean and SEM across genes in each bin. Note that the scales on the y-axis are different between species.

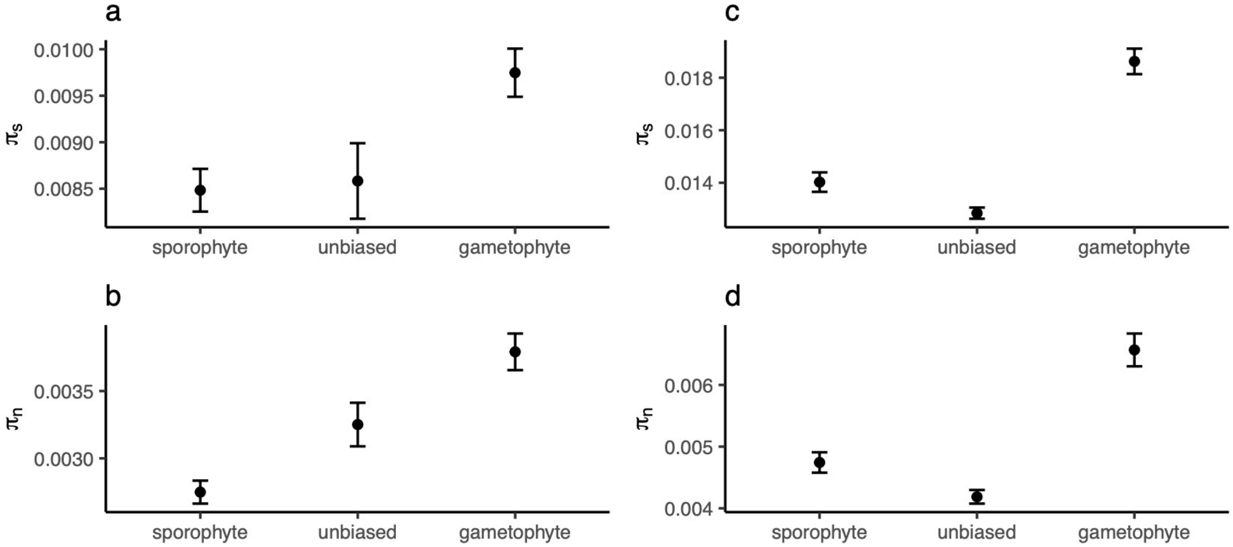


Figure S4. Mean nucleotide diversity of gametophyte-specific, sporophyte-specific, unbiased genes in *R. hastatulus* (a-b) and *C. purpureus* (c-d). The numbers of genes in each category are the same as Figure 2. Error bars represent mean and SEM across genes in each category. Note that the scales on the y-axis are different between species.

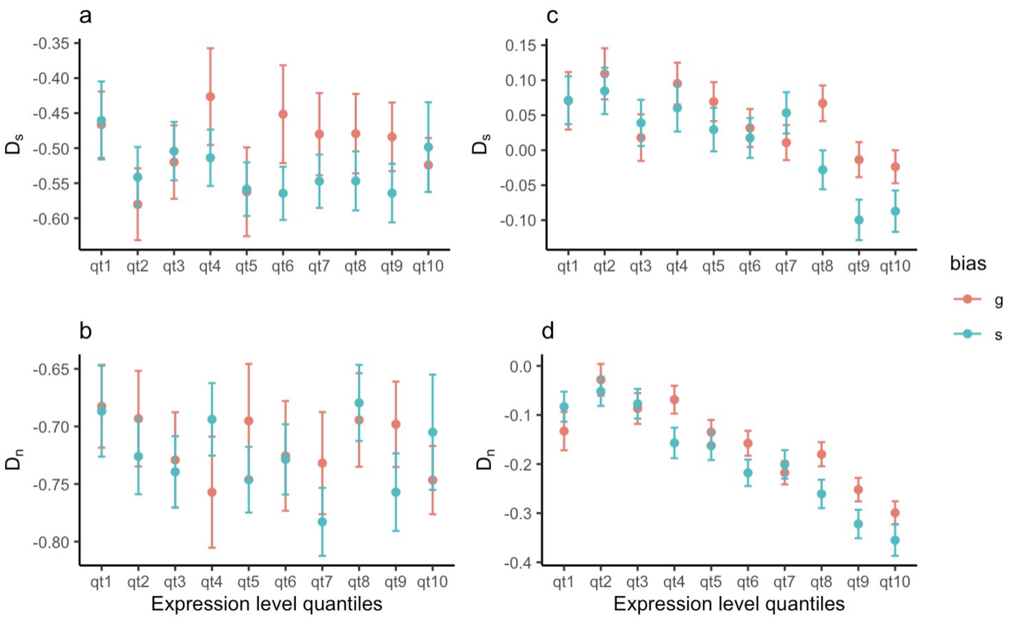


Figure S5. Effect of expression bias between life stages and expression level on Tajima's *D*. Number of genes in each bin: 676-843 ($D_s$), 1,150-1,339 ($D_n$) for *R. hastatulus* (a, b); 1,265-2,250 ($D_s$), 1,456-2,090 ($D_n$) for *C. purpureus* (c, d). qt1: lowest expression level, qt10: highest expression level. Error bars represent mean and SEM in each bin. Note that the scales on the y-axis are different between species.

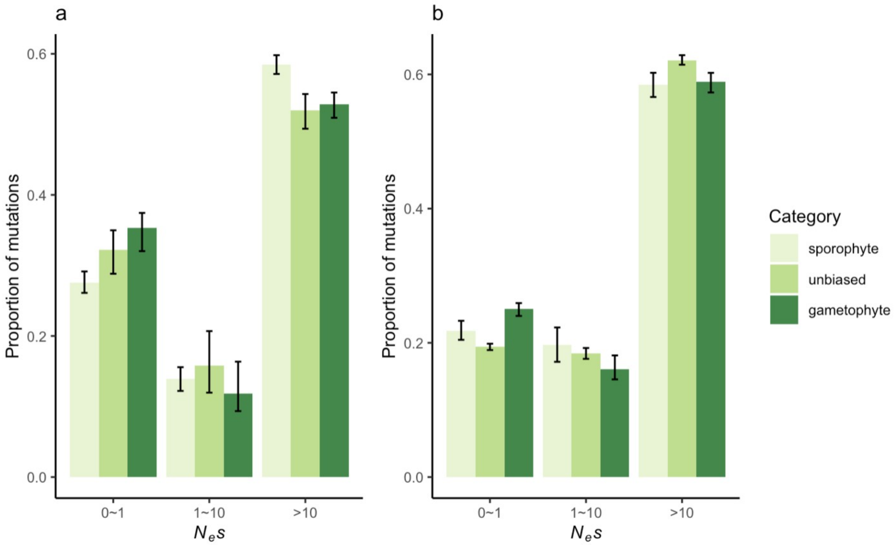


Figure S6. Distribution of fitness effects of life stage-specific genes in *R. hastatulus* (a) and *C. purpureus* (b). Number of genes in each category: 1771 (g), 1771(s) for *R. hastatulus* 1514 (g), 1656 (s) for *C. purpureus*. The mean and 95% confidence intervals are based on 200 bootstraps of the original dataset.

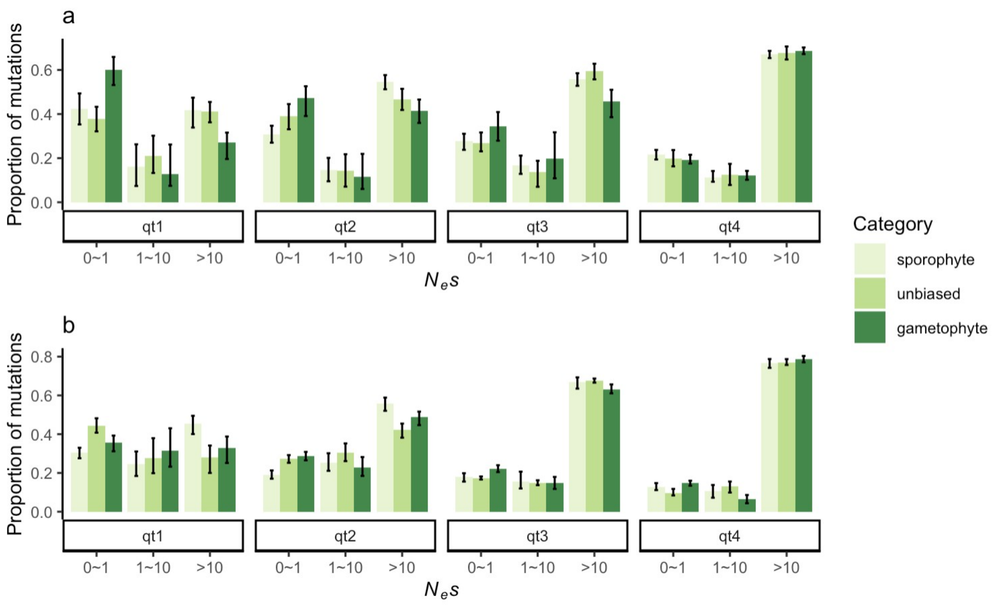


Figure S7. Distribution of fitness effects of life stage-specific genes across expression level quantiles in *R. hastatulus* (a) and *C. purpureus* (b). Number of genes in each quantile of expression level: 322, 407, 457, 525 (s) and 570, 254, 180, 707 (g) for *R. hastatulus*; 873, 429, 197, 157 (s) and 489, 454, 272, 298 (g) for *C. purpureus*. The mean and 95% confidence intervals are based on 200 bootstraps of the original dataset.

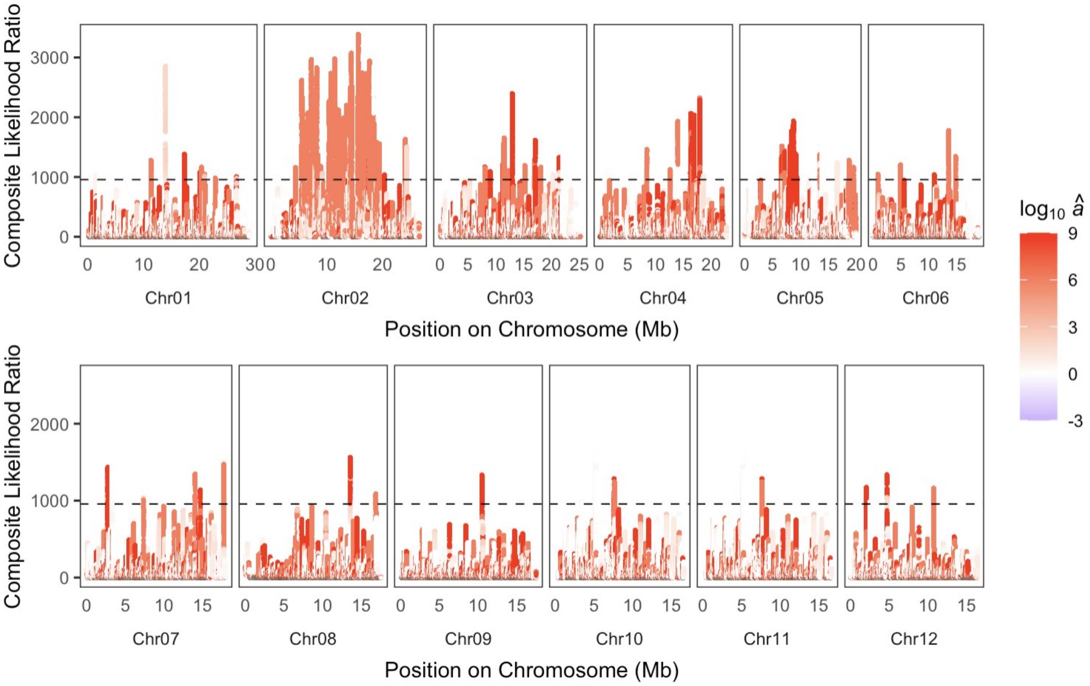


Figure S8. Genome-wide scan for balancing selection in *C. purpureus*. Black dashed lines represent the cutoff for the signal of balancing selection. $\hat{a}$: estimated dispersion parameter, a positive $\log_{10}\hat{a}$ value suggests balancing selection.

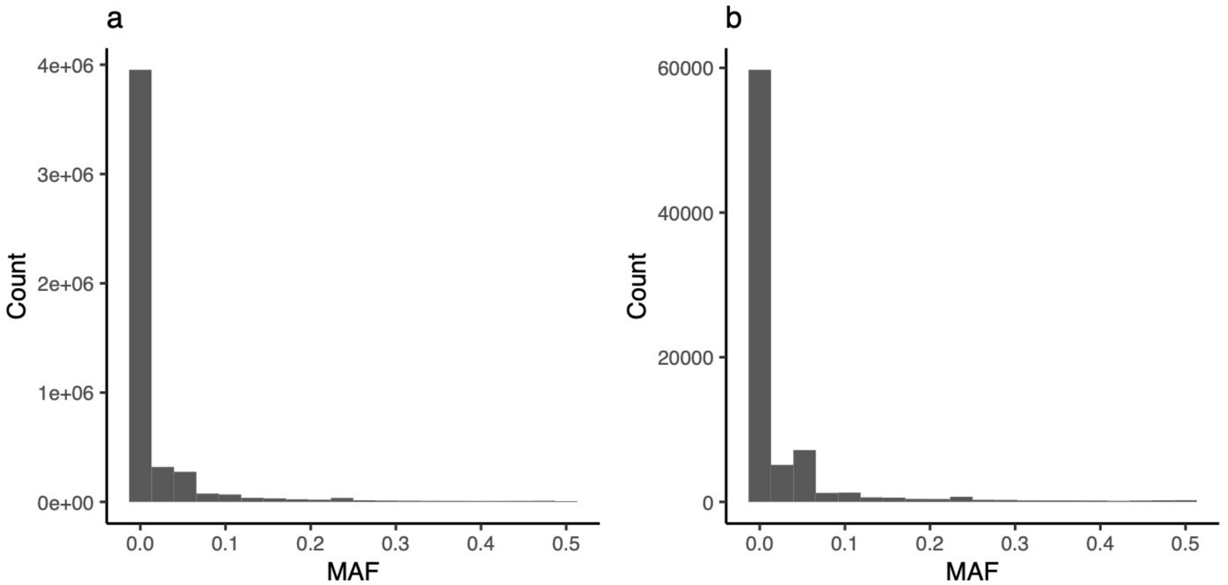


Figure S9. Distribution of minor allele frequency (MAF) of all sites tested (a) and candidate sites under balancing selection (b) in *R. hastatulus*. Mean MAF: 0.02 (a), 0.025 (b); median MAF: 0 (a), 0 (b); proportion of sites with MAF > 0.3: 0.016 (a), 0.021 (b).

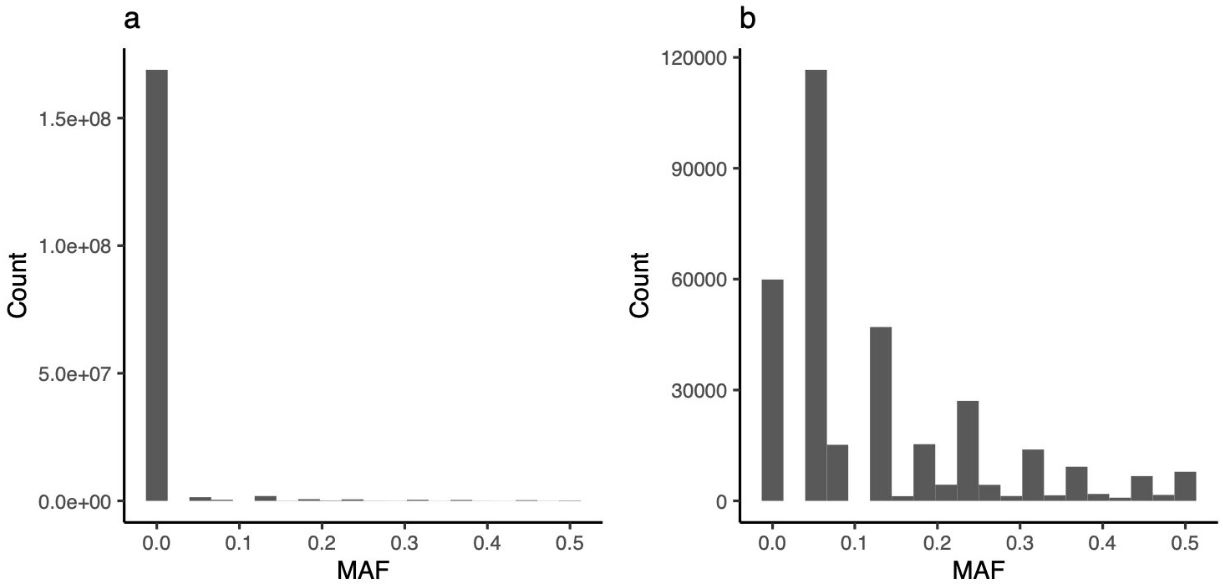


Figure S10. Distribution of minor allele frequency (MAF) of all sites tested (a) and candidate sites under balancing selection (b) in *C. purpureus*. Mean MAF: 0.0072 (a), 0.13 (b); median MAF: 0 (a), 0.063 (b); proportion of sites with MAF > 0.3: 0.008 (a), 0.13 (b).

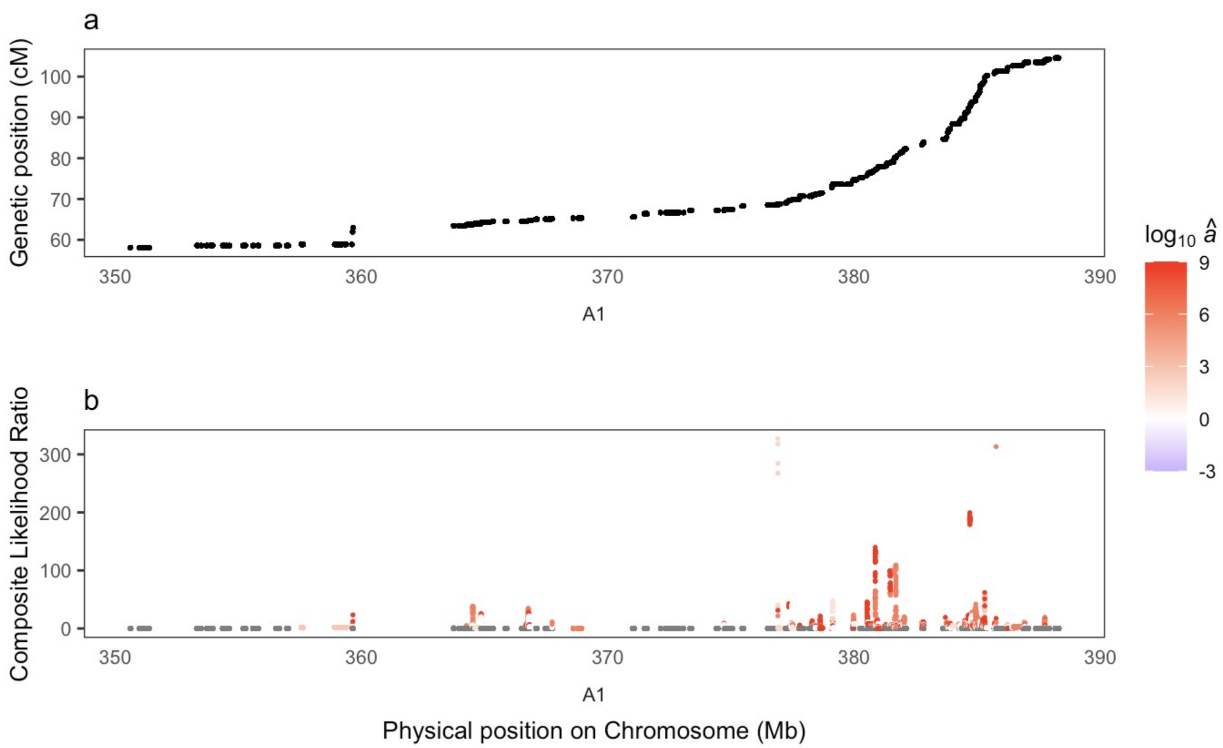


Figure S11. Scans of balancing selection in *R. hastatulus* in a 40 Mb window at the end of A1. y-axis: genetic positions of sites being tested (a), composite likelihood ratio (b).
